# Pivoting of microtubules driven by minus end directed motors leads to their alignment to form an interpolar bundle

**DOI:** 10.1101/347831

**Authors:** Lora Winters, Ivana Ban, Marcel Prelogović, Nenad Pavin, Iva M. Tolić

**Author notes:** These authors contributed equally to this work (Lora Winters, Ivana Ban, Marcel Prelogović). Correspondence (I.M.T.) or (N.P.).

## Abstract

At the beginning of mitosis, the cell forms a spindle made of microtubules and associated proteins to segregate chromosomes. An important part of spindle architecture is a set of antiparallel microtubule bundles connecting the two spindle poles. A key question is how microtubules extending at arbitrary angles form an antiparallel interpolar bundle. Here we show that microtubules meet at an oblique angle and subsequently rotate into antiparallel alignment. By combining experiments with theory, we show that microtubules from each pole search for those from the opposite pole by performing random angular movement. Upon contact of two microtubules, they slide sideways along each other towards the minus end, which we interpret as the action of minus end directed Cut7/kinesin-5 motors. In conclusion, random rotational motion helps microtubules from the opposite poles to find each other and subsequent accumulation of motors allows them to generate forces that drive interpolar bundle formation.

---

During cell division, the genetic material is divided into two equal parts by the mitotic spindle. This complex dynamic micro-machine is made of microtubules (MTs) emanating from the spindle poles, chromosomes and a variety of accessory proteins^1,2^. Some MTs extending from the spindle pole are bound to kinetochores on the chromosome, whereas others are bound to MTs extending from the opposite pole, in an antiparallel configuration known as interpolar or overlap bundles^3–6^. These bundles interact laterally with kinetochore MTs and regulate the forces acting on chromosomes and spindle poles^7–12^.

MTs within interpolar bundles are cross-linked by specific proteins, which can be divided into three classes: (i) motors that slide the MTs and thus the spindle poles apart by walking along the MTs away from the pole, i.e., towards the plus end of the MT, such as kinesin-5 motors Cut7/Cin8/Eg5/KIF11 (ref.^13–15^); (ii) motors that pull the poles together by walking along the MTs towards the pole, i.e., towards the minus end of the MT, such as kinesin-14 motors Ncd/HSET/KifC1 (ref.^16,17^); (iii) proteins that cross-link MTs without walking along the MTs, such as Ase1/PRC1 (ref.^18^). Remarkably, *in vitro* studies have shown that kinesin-5 motors can also move towards the minus end of the MTs, when walking on a single MT or in a non-crowded environment on antiparallel MTs^19–23^. Likewise, kinesin-14 motors can reverse the direction of movement under a low external force^24^. Stability of antiparallel bundles for combinations of motors and crosslinkers has been explored theoretically^25,26^.

While the already formed antiparallel bundles in metaphase have been to a large extent described, little is known about how these highly organized structures are formed during prometaphase. The reason is that this dynamic process is not accessible by current experimental techniques due to a high number of MTs extending from the spindle poles in prometaphase in higher eukaryotic cells^27^, which may form antiparallel bundles. In yeast cells, which have a small number of MTs and a rod-shaped spindle^4,6^, the study of the antiparallel bundle formation in living cells is challenging because the spindle poles are next to each other at the onset of prometaphase. Yet, the advantage of yeasts as experimental systems is that their spindles consist of only one antiparallel bundle. Electron tomography on early spindles in yeast showed MTs interacting at oblique angles^28^, suggesting that such interactions may be an intermediate step during the formation of antiparallel bundles.

Alignment of MTs into an antiparallel configuration may be achieved by rotation of MTs that initially extend at an oblique angle. Indeed, live-cell imaging in fission yeast showed that MTs change their angle as they rotate (i.e., pivot) around the spindle pole^29^. Eventually these MTs join the spindle, with help from Ase1 crosslinkers^30^. Experiments on budding yeast showed that cells lacking kinesin-14 motors have more MTs extending at an oblique angle with respect to the spindle and fewer antiparallel MTs than wild-type cells do^31^. Based on this finding, the authors hypothesized that minus end directed motors align the MTs, which was verified by computer simulations^31^. Spindle assembly starting from a monopole has been explored by extensive computer simulations including motors of different directionalities and passive crosslinkers^32^. Experiments together with simulations showed that spindles can form even in the absence of motors, in which case MT growth and Ase1 crosslinkers play an important role^33,34^. Even though the formation of antiparallel bundles has been explored theoretically, a key question remains of how this process occurs *in vivo.*

## RESULTS

### Assay for spindle reassembly in fission yeast

At the onset of mitosis in the fission yeast *Schizosaccharomyces pombe*, the two spindle pole bodies (SPBs) are embedded in the nuclear envelope, which remains intact during mitosis mitosis^35^. The SPBs nucleate polar MTs with minus ends at the SPBs and the plus ends in the nucleoplasm 4,36. MTs extending from the opposite SPBs interact and form an antiparallel interpolar bundle, and together with MTs that bind to kinetochores assemble the spindle. The interactions between antiparallel MTs occur when the SPBs are next to each other^37^, making it difficult to study the dynamics of this process. To increase the distance between the SPBs, we used a spindle reassembly assay, in which we disassembled the spindle by exposing the cells in metaphase to cold temperature (1°C, Fig. 1a, Supplementary Fig. S1a), adapting the approach that was previously used to study kinetochore capture^29,38^. SPBs were visualized by Sid4-GFP and MTs by GFP-tubulin. When the temperature was increased to permissive temperature (24°C), the SPBs were more than 1 μm apart in 61±5% of cells (n=84; results are mean±s.e.m. unless otherwise stated) (Supplementary Fig. S1b). Thus, this assay allowed us to investigate the process of antiparallel bundle formation, i.e., spindle reassembly.

### Antiparallel microtubule bundles are formed in two steps

After the cold treatment was ended and the cells returned to permissive temperature, 75±5% (65 out of 87) spindles reassembled within 10 minutes, which is a typical duration of prophase and metaphase in unperturbed mitosis^39^. Shortly after the return to permissive temperature, MTs started growing from each of the two SPBs (defined as time 0, Fig. 1b). MTs did not extend in a defined direction, but instead pivoted around the SPB (Fig. 1b, 0:00-1:22), in agreement with our previous observations^29^. Eventually a MT extending from one SPB came into contact with a MT from the other SPB (Fig. 1b, 1:22). At the time of initial contact, MTs were typically not aligned in an antiparallel manner, but interacted at an oblique angle (Fig. 1b, 1:22). In 21 out of 31 cells that reassembled their spindles and had the SPBs separated by more than 1 μm, MTs interacted at an oblique angle, in 6 they met at the pole-pole axis, and in 4 one MT grew directly to the opposite pole. Following the initial contact, MTs rotated into antiparallel alignment (Fig. 1b, 1:22-1:46, Supplementary Movie 1; note that not every apparent contact led to alignment). Thus, formation of an antiparallel bundle occurs in two steps: (i) MT growth and random rotation before their contact, which we refer to as search, and (ii) directed rotation of MTs towards an antiparallel configuration, which we term aligning (Fig. 1c).

**Figure 1.**
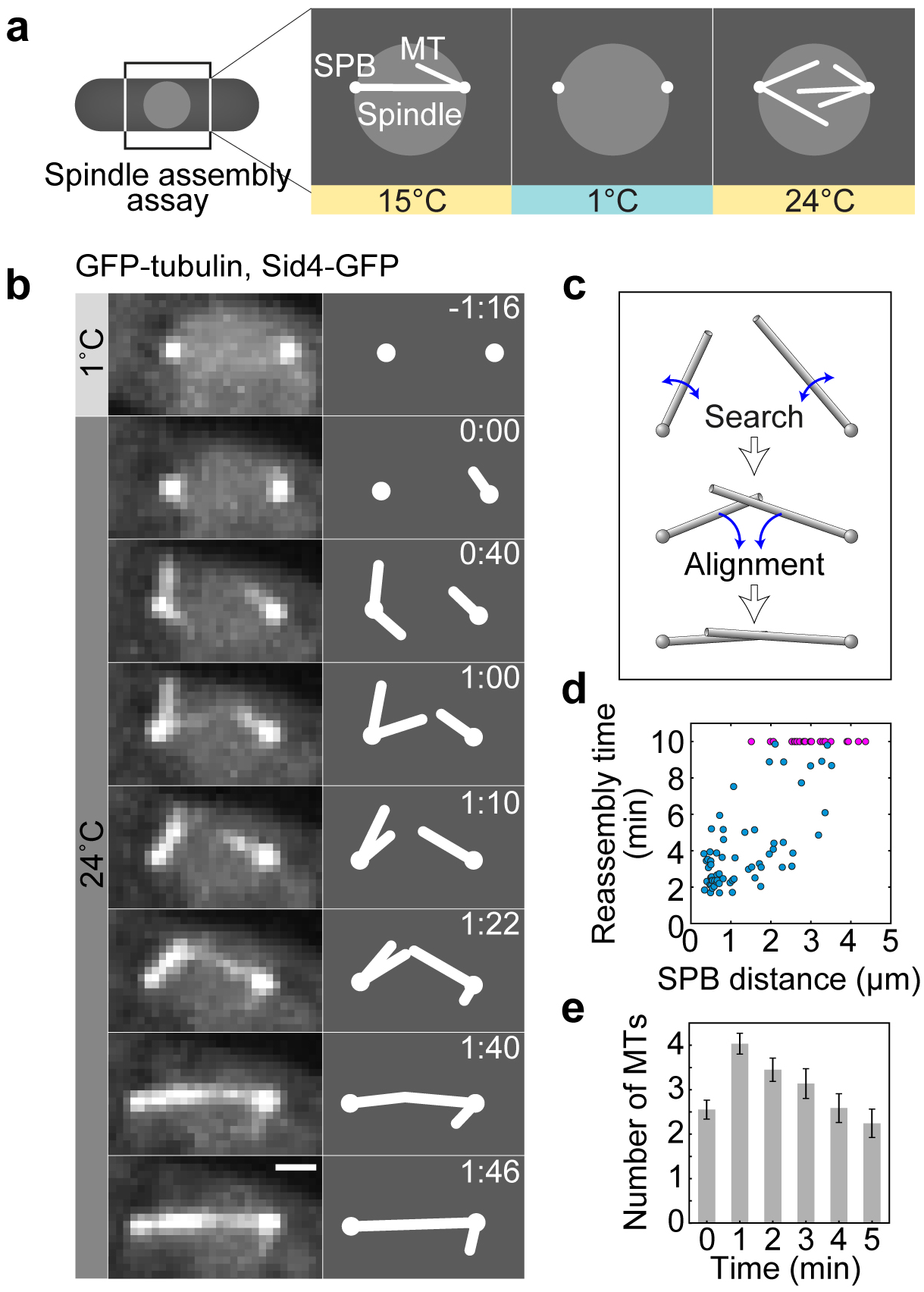
Spindles reassemble by rotational movements of MTs. (**a**) Spindle reassembly assay. Mitotic cells were cooled to 1°C to depolymerize MTs (Methods). Once the temperature was increased to 24°C, MTs grew from the SPBs and reassembled the spindle. (**b**) Time-lapse images of spindle reassembly in a wild-type cell expressing GFP-tubulin and Sid4-GFP (strain KI061). Images are maximum-intensity projections, time is given in min:s, scale bar, 1 μm. Corresponding schemes are shown to the right. (**c**) Scheme showing how the formation of an antiparallel MT bundle occurs in two steps. SPBs are represented as spheres and MTs as rods. (**d**) Reassembly time, defined as the time from the onset of MT growth until the formation of the antiparallel MT bundle (spindle) between the SPBs, as a function of the distance between the SPBs at the onset of MT growth. n=87 cells; pink data points denote cells in which the spindle was not reassembled within 10 minutes. (**e**) Number of polar MTs per cell during the first 5 minutes following the onset of MT growth, n=28 cells, error bars, s.e.m.

### Quantification of spindle reassembly

To quantify the kinetics of spindle reassembly, we measured the spindle reassembly time, defined as the time needed for the formation of an antiparallel MT bundle between the SPBs, which includes both steps of this process. The average reassembly time was 7.3±0.9 min (n=87 cells). The cells with a larger initial distance between the SPBs took a longer time to reassemble the spindle or did not reassemble the spindle within 10 minutes (Fig. 1d).

To describe the first step of bundle formation, in which MT contact is established, we quantify polar MTs and their movement. During the process of spindle reassembly, the average number of polar MTs per cell increased from 0 to 2.5 during the first minute and to 4 in the second minute (n=28 cells, Fig. 1e). Afterwards, the number of MTs decreased. MTs typically reached a length of 1.5 μm (Supplementary Fig. S1c) and their angular diffusion coefficient was 4.5 degrees^2^/s (Supplementary Fig. S1d), in agreement with our previous measurements^29^. Note that the structure that appears as a polar MT may be a bundle of a few MTs; thus our measurements correspond to the bundle.

### Microtubules slide sideways towards each other’s minus end during the process of alignment

In the second step of bundle formation MTs rotate into antiparallel alignment by sliding sideways along each other towards the SPB (Fig. 2a). To quantify the geometry of this system over time, we first measured the angle between the MTs, α (Fig. 2b), starting 10 seconds before the antiparallel bundle was formed. We found that the angle increased towards 180 degrees, which represents the bundled configuration (Fig. 2c). During the subsequent 4 seconds the angle remained constant (Fig. 2c).

Next, we measured the total contour length of the MTs that formed an antiparallel bundle, defined as a segmented line starting at one SPB, passing through the contact point between the MTs, and ending at the other SPB (Fig. 2b, Supplementary Fig. S1e; Methods). We found that the total contour length of the MTs decreased during the rotation of the MTs into antiparallel alignment, and remained constant afterwards (Fig. 2d). The contour length decreased in a roughly linear manner, which allowed us to introduce the measure termed contour velocity, defined as the velocity at which the contour length changes. We measured a contour velocity of −18±2 nm/s (n=14 cells). Contrary to the contour length, the distance between the SPBs was constant both during the alignment and afterwards (Fig. 2e). Given that MTs do not undergo poleward flux in fission yeast^40^, our results reveal that during the alignment the contact point between the MTs moves in a directed manner towards the minus end of each MT, which is at the SPB. Thus, the alignment may be driven by minus end directed motors. The velocity at which the motors slide the MTs with respect to each other equals the velocity of change of the contour length.

**Figure 2.**
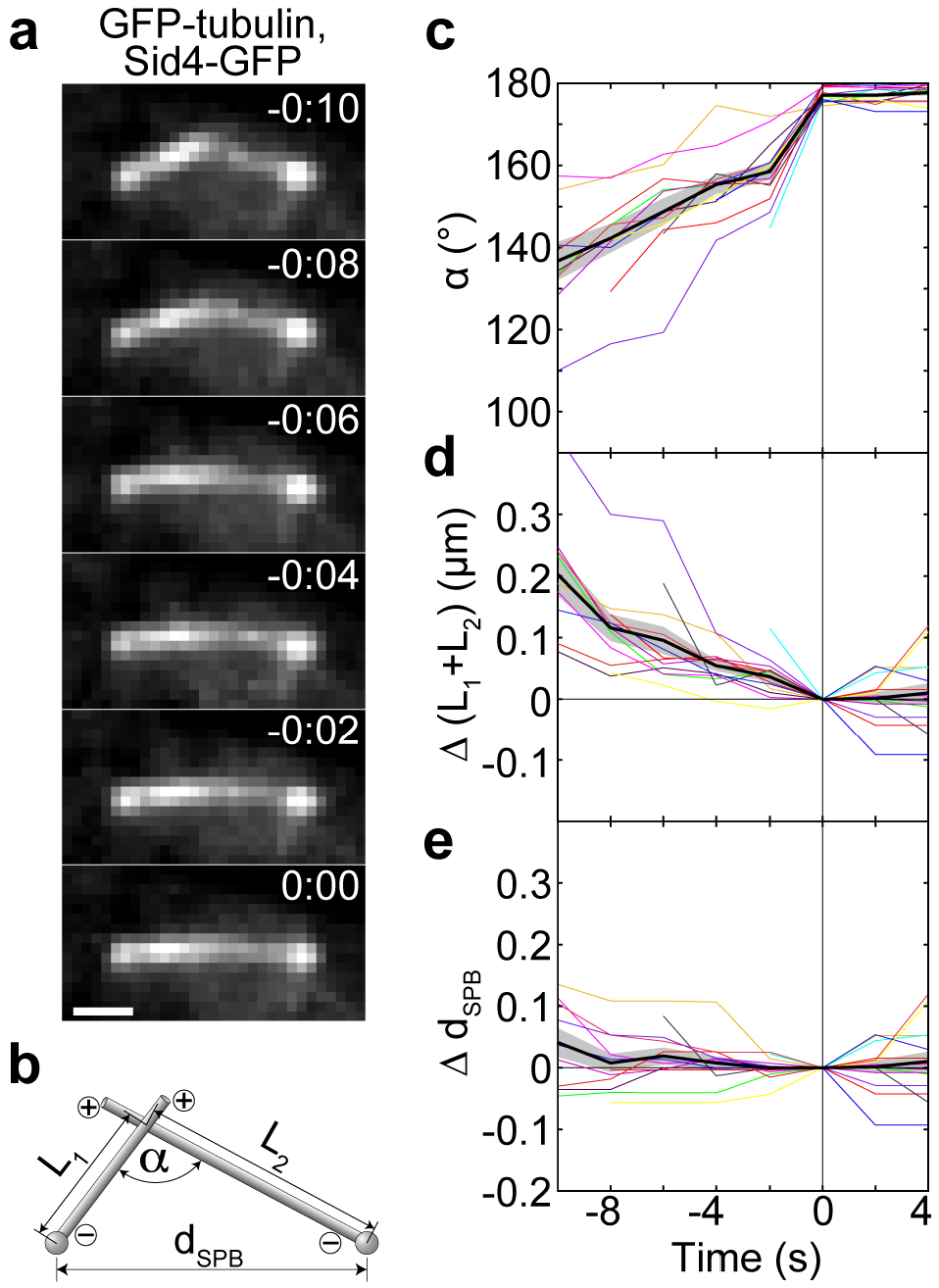
MTs slide sideways towards each other’s minus end during alignment. (**a**) Time-lapse images of a wild-type cell expressing GFP-tubulin and Sid4-GFP (strain KI061) during the formation of an antiparallel bundle. Images are maximum-intensity projections, time is given in min:s, scale bar, 1 μm. (**b**) Measurement of the angle between the MTs, *α*, the contour length of MTs, *L_1_ + L_2_*, and the distance between the SPBs, *d_SPB_*, during the formation of an antiparallel bundle. SPBs are represented as spheres and MT s as rods; plus and minus signs designate the respective ends of MTs. (**c**) Angle between MTs, *α*, as a function of time. (**d**) Contour length difference as a function of time. The contour length difference is defined as the difference between the contour length, *L_1_* + *L_2_*, at a given time and the contour length at *t* = 0, Δ(*L*_1_ + *L*_2_) = *L*_1_ + *L*_2_ − (*L*_1_ + *L*_2_)|_*t*=0_. (**e**) The difference of distances between the SPBs as a function of time, defined as the difference between the SPB distance, *d*_SPB_, at a given time and the distance at *t*=0, Δ*d*_SPB_ = *d*_SPB_ − *d*_SPB_|_*t*=0_. In panels (**c**)-(**e**), the same strain as in (a) was used; time 0 is the time when the antiparallel bundle was formed; n=14 cells; individual cells (colored lines), mean value (black line), and s.e.m. (shaded area) are shown.

### Cut 7 (kinesin-5) is important for spindle reassembly

We explored the role of motor proteins and a non-motor MT crosslinker in spindle reassembly. First we focused on Cut7, a kinesin-5 family member, because it is essential for spindle formation^13,41^, whereas spindles are able to assemble without any of the other 8 kinesins in *S. pombe*^42–47^, dynein^48^ and the non-motor crosslinker Ase1/PRC1 (ref.^49,50^). We used a temperature-sensitive *cut7.24*^ts^ mutant in our spindle reassembly assay and set the final temperature to 37°C to abrogate Cut7 activity. Similar temperatures do not disrupt spindle assembly and completion of mitosis in wild-type cells^39^. We estimate that Cut7 was inactivated within 3 minutes, given that it took about a minute to raise the temperature from 1°C to 37°C and the mutation response time was estimated to be 2 minutes^47^. We found that only 5±5% (1 out of 19) spindles in *cut7.24*^ts^ cells reassembled at times longer than 3 minutes (Fig. 3a). In the remaining cells the MTs extending from the two SPBs did not form an antiparallel bundle, often crossing each other in an X-shaped conformation (Fig. 3b, Supplementary Movie 2). The fraction of reassembled spindles was smaller than the fraction of reassembled spindles in wild type at times longer than 3 minutes (63±6%, or 38 out of 60, calculated from data in Fig. 1d). To exclude the possibility that this difference is due to different distances between the SPBs, we took into account only the cells in which this distance was in the range 1-5 μm, and found that 14±8% (3 out of 21) spindles in the *cut7.24*^ts^ mutant reassembled, whereas in wild type this fraction was 58±7% (31 out of 53). Additional comparisons are shown in Supplementary Fig. S1f. We conclude that Cut7 is important for the formation of an antiparallel bundle in the spindle reassembly assay.

We also applied the spindle reassembly assay to the mutants lacking the proteins that have been shown to regulate spindle length in metaphase: kinesin-8 motor protein Klp5 (ref.^51,52^), kinesin-14 motors Pkl1 and Klp2 (ref.^42,43^), and the crosslinker Ase1 (ref.^49,50^). We found that spindles were able to reassemble in the absence of these proteins (Supplementary Fig. S1g, h). In all the studied mutants, the reassembly time increased with an increase in the distance between the SPBs, similarly to wild type (Supplementary Fig. S1i).

To compare *cut7.24*^ts^ cells with the other mutants and wild type, we analyzed only the cells that reassembled spindles at times longer than 3 minutes or did not reassemble. Whereas in wild type and all the mutants except *cut7.24*^ts^ more than 50% of the spindles reassembled, in *cut7.24*^ts^ this fraction was only 5% (Supplementary Fig. S1j). Taken together, our experiments on a set of mutants suggest that Cut7 has a function in the transformation of oblique MT contacts into antiparallel bundles to reassemble the spindle.

### Cut7 is found at the contact site between microtubules during microtubule rotation into antiparallel alignment

To explore the localization of Cut7 and thus the potential sites where it may exert forces that align the MTs from the opposite SPBs into antiparallel configuration, we used cells expressing Cut7-3GFP as well as mCherry-tubulin and Sid4-mCherry in our assay (Fig. 3c, Supplementary Fig. S2a-c, Supplementary Movie 3). During cold treatment, Cut7-3GFP showed a diffuse signal in the nucleus (Supplementary Movie 3). When the temperature was increased and MTs started to nucleate from the SPBs, Cut7 appeared close to the SPBs (n=50 out of 50 cells; Fig. 3c, 0:10). We first analyzed the cells in which the SPBs were separated by more than 1 μm after the cold treatment (n=19 out of 50 cells). We found that MTs pivoted around the SPBs and eventually formed a bundle connecting the SPBs (n=15 out of 19 cells; Fig. 3c, 0:100:35), in agreement with our results shown in Fig. 1b. Interestingly, when MTs extending from the two SPBs established contact, Cut7 was found at the contact site (n=8 out of 8 cells in which the initial MT contact was clearly visible; Fig. 3c, 0:25; in the remaining 7 cells the initial MT contact site was unclear). Intensity profiles of Cut7-3GFP signal along the contour length of MTs show that Cut7 appeared at the contact point and was present at this point during the process of MT alignment (Fig. 3d, 0:25-0:35). When the angle between the MTs changed from an oblique to the straight angle, Cut7 distribution changed from a spot to a broader distribution along the spindle, resulting in several Cut7 streaks along the spindle (n=14 out of 15 cells; Supplementary Movie 3). Similar examples are shown in Supplementary Fig. S2a,b.

In the cells in which the SPBs were separated by less than 1 μm after the cold treatment (n=31 out of 50 cells), Cut7 was found at the SPBs upon temperature increase (n=31 out of 31 cells). The spindles reassembled (n=30 out of 31 cells), but it was not possible to observe the initial contact between the MTs extending from the opposite SPBs and the distribution of Cut7 at that time due to the short distance between the SPBs (Supplementary Fig. S2c). Note that the kinetics of spindle reassembly in the cells expressing Cut7-3GFP (Supplementary Fig. S2d) was similar to that in wild-type cells shown in Fig. 1d, suggesting that labeling of Cut7 did not perturb this process.

Based on our experiments in which Cut7 and MTs were visualized, we conclude that Cut7 near the SPBs cannot contribute to the alignment of MTs extending from the opposite SPBs and interacting at an oblique angle, because all MTs extend from the same SPB in that region. We speculate that Cut7 found at the site of MT interaction may exert forces that align the MTs into antiparallel configuration.

**Figure 3.**
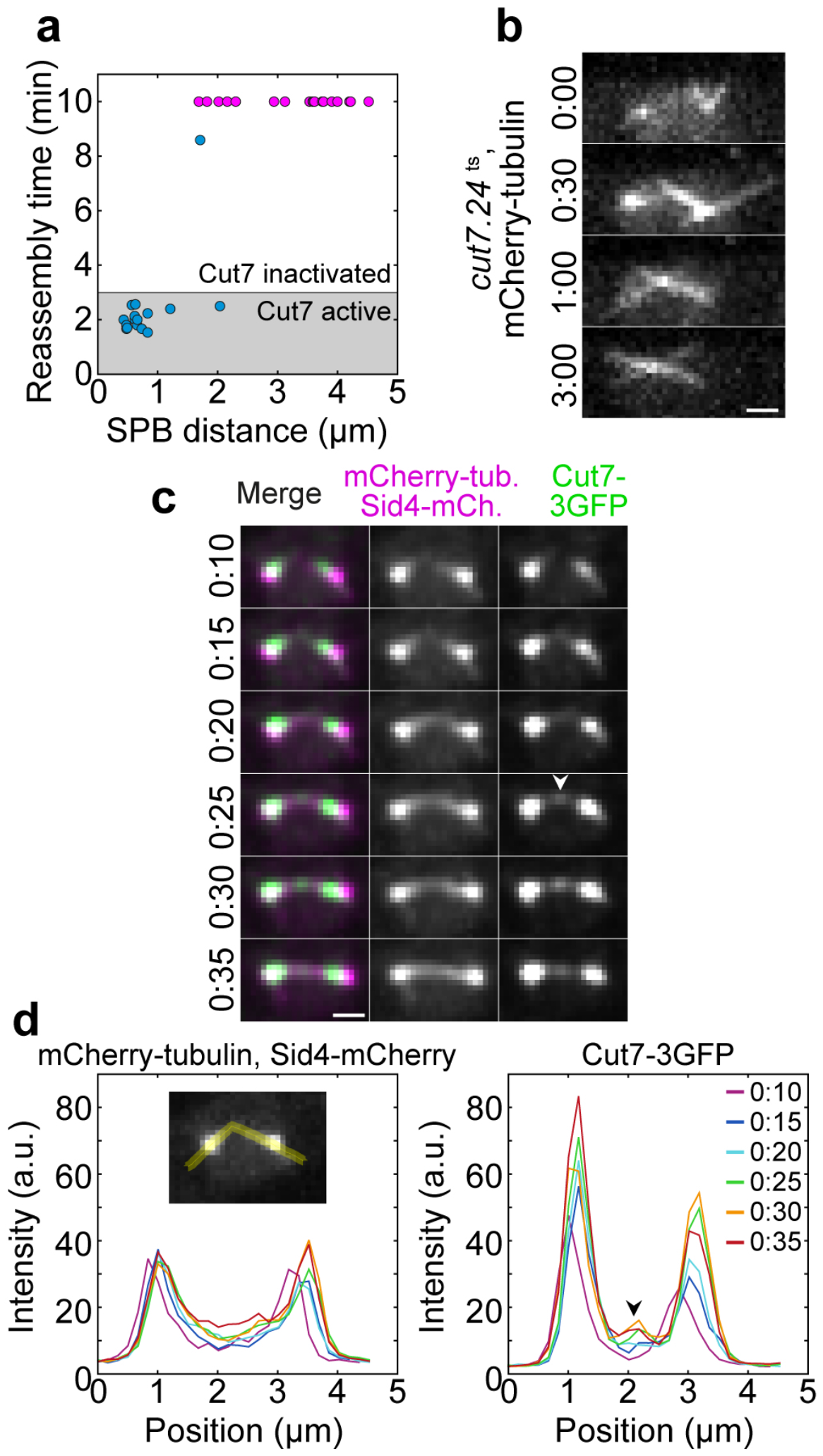
Cut7-GFP accumulates at the initial contact points of MTs. (**a**) Reassembly time as a function of the distance between the SPBs for cut7.24^ts^ cells (strain CF.391) at non-permissive temperature (37°C), n=34 cells. Pink data points denote cells in which the spindle was not reassembled within 10 minutes. (**b**) Time-lapse images of a cut7.24^ts^ cell expressing mCherry-tubulin (strain CF.391) at non-permissive temperature (37°C), in which the spindle did not reassemble. (**c**) Spindle reassembly in a cell expressing Cut7-3GFP (green), mCherry-tubulin (magenta), and Sid4-mCherry (magenta; strain LW042). Merged time-lapse images (left column) and separate channels (central and right column, both in gray scale) are shown. Note the accumulation of Cut7 at the site of MT contact (arrowhead). (**d**) Signal intensity profiles of mCherry-tubulin (left) and Cut7-3GFP (right) measured along the MT contour extended into the cytoplasm (see example in the inset on the left), at times noted in the legend on the right. The arrowhead marks the peak of Cut7-3GFP signal in the MT contact region. The intensity profiles were measured on the images shown in panel (**c**). In (**b**) and (**c**), images are maximum-intensity projections, time is given in min:s; scale bars, 1 μm.

### Theoretical model

To explore how the MTs, which extend in arbitrary directions, become aligned into an antiparallel bundle connecting the spindle poles, we introduce a simple physical model (Fig. 4a and Methods). The central idea of our theoretical approach is that MTs perform rotational movement around the spindle pole, allowing them to explore the space as they search for the MTs extending from the opposite pole and to establish a configuration required for spindle assembly. In our model, two types of forces drive the rotational movement of MTs: forces generated by motor proteins and thermal forces. The forces generated by motors appear when MTs get into close proximity allowing the motors to attach in this region and thus crosslink the MTs. A motor is described as an elastic spring, whose two ends can move along two MTs. The motors move towards the MT minus end, which is at the spindle pole, generating a directed force on the MTs that rotates them towards the pole-pole axis. A motor is considered as a force generator, whose velocity decreases under load. In contrast to motor-generated forces, thermal forces are random and always present irrespective of the distance between the MTs. To keep the model simple, we consider straight MTs of a constant length extending from each spindle pole. MTs are pinned at one end at the nuclear envelope of a spherical shape. We use this model to calculate the dynamics of antiparallel bundle formation.

We solved the model numerically to obtain the time course for MT orientations and the number of attached motors. MT orientations are parametrized by angular coordinates, the polar and the azimuthal angle (Fig. 4a). For parameters given in Table 1 and discussed in the Methods section, solutions of the model show that MTs initially preform random angular movement (dark gray region in Fig. 4b, top). This movement of MTs is predominantly driven by thermal forces, as there are no motors attached to them (Fig. 4b, middle) because the MTs are not yet in contact (Fig. 4b, bottom). This random movement of MTs corresponds to the random movement of MTs observed in experiments (Fig. 1b, 0:00-1:22), which is typical for the search step. Our calculations show that this random movement ends when the MTs come close enough to each other so that motors can attach (Fig. 4b, bottom). Subsequently, polar angles change in a directed manner towards the antiparallel configuration (light gray region in Fig. 4b, top). This directed movement is the result of the accumulation of motors that generate forces (Fig. 4b, middle). The movement stops when the polar angles approach 0° and 180° for the first and the second MT, respectively (end of the light gray region in Fig. 4b, top), thereby forming a stable antiparallel bundle. This directed movement of MTs and the accompanying accumulation of motors correspond to the experimental observations that characterize the alignment step (Figs. 2 and 3). The entire time course of the MTs and the behavior of motors corresponding to Fig. 4b is illustrated in Supplementary Movie 4, and the still frames from the animation representing the search, aligning and bundled state are shown in Fig. 4c, top, middle and bottom, respectively.

**Figure 4.**
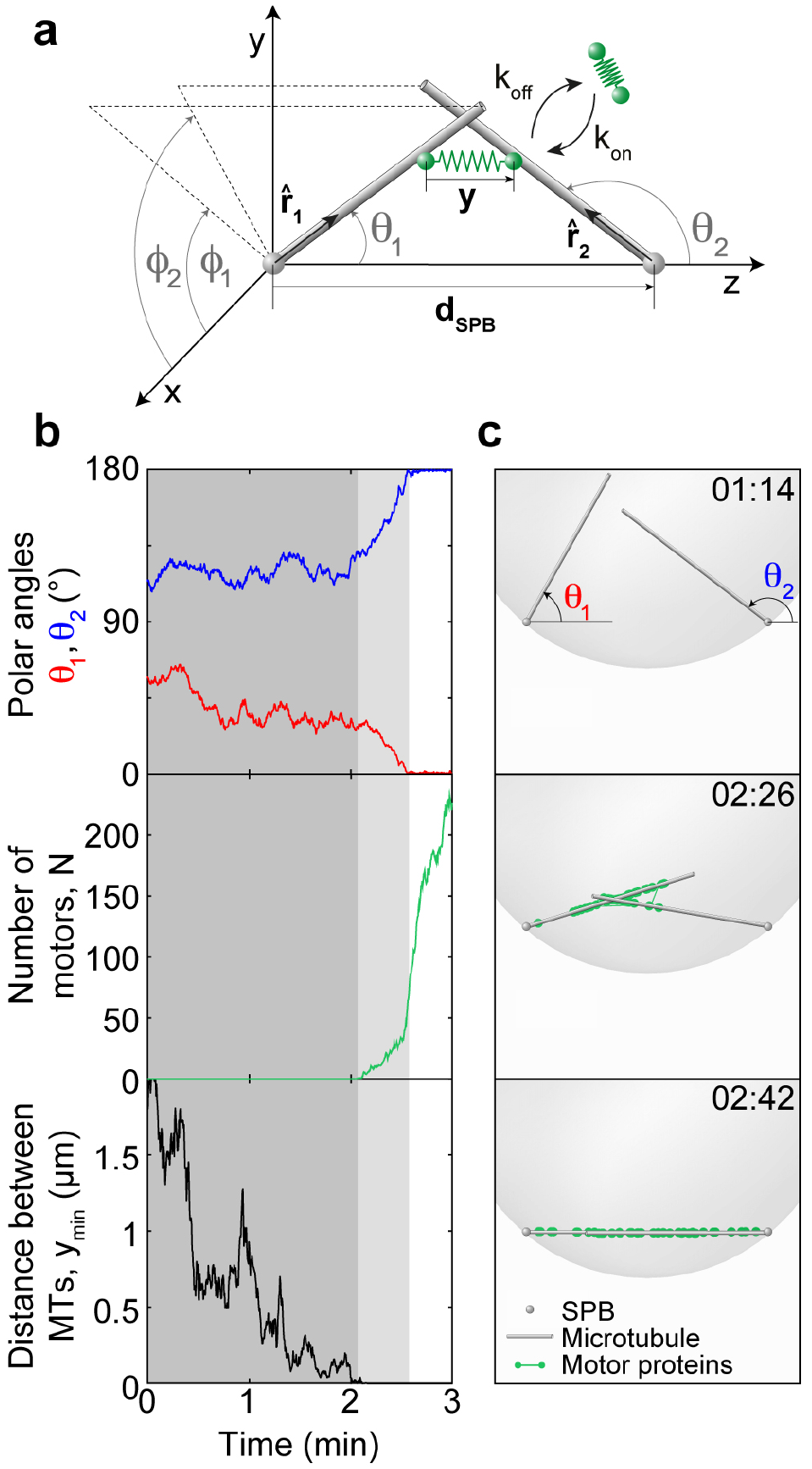
Theoretical model and solutions for MT dynamics. (**a**) Scheme of the model. Each MT (gray rod) is freely joint to its respective SPB (gray sphere). Orientations of two MTs are represented with unit vectors **r̂**_1_and **r̂**_2_ respectively, while the SPBs are at fixed points separated by the distance **d**_SPB_. Motor proteins (green springs) can attach to and detach from MTs with rates *k*_on_ and *k*_off_, respectively, and when attached, their elongation is *y.* In the Cartesian coordinates, the SPBs are at points (0,0,0) and (0,0, *d*_SPB_). The MT orientations are described by the polar angles *θ*_1_ and *θ*_2_ and by the azimuthal angles and *ø*_1_ and *ø*_2_ for the first and the second MT respectively. (**b**) A sample path representing a bundling event. Top graph represents the polar angles, which are denoted with a blue and a red line for the first and second MT respectively. In the antiparallel configuration, the first MT has the polar angle *θ*_1_ = 0° and the second *θ*_2_ = 180° (see (**a**) for parametrization). The middle graph shows the number of attached motors and the bottom graph shows the distance between the two closest points on the MTs. The shaded regions represent the search and the aligning phase (dark and light gray respectively) and the non-shaded region represents the bundled state. Simulation is performed with parameter values *R*_1,2_ = 1.5 *μm*, *d*_SPB_ = 2 *μm*, *n*_MT_ = 2, and other values from Table 1. (**c**) Illustrations of search (top), aligning (middle) and bundled state (bottom) of MTs (gray rods). To illustrate the motor distribution, the position of each motor (green) is randomly generated using a normal distribution around their mean position with the steady state variance, which is calculated from the MT orientations. The small gray spheres represent the SPBs and the large translucent gray sphere represents the nuclear envelope. The images are taken from Supplementary Movie 4, which is produced using the same data as in (**b**). Time is given in min:s.

**Table 1.** Parameters used in the model. The choice of parameters values is described in Methods.

| Parameter |  | Value | Source |
| --- | --- | --- | --- |
| $v_0$ | Motor velocity | $-0.01 \mu\text{m/s}$ | Estimated from contour velocity |
| $D_v$ | Motor velocity fluctuation | $5 \times 10^{-4} \mu\text{m}^2/\text{s}$ | Ref.<sup>53</sup> |
| $f_0$ | Motor stall force | $-1.5 \text{ pN}$ | Ref.<sup>54</sup> |
| $k_{\text{on}} c_{\text{nuc}}$ | Motor concentration parameter | $1.3 \text{ s}^{-1} \mu\text{m}^{-1}$ | Estimated |
| $k_{\text{off}}$ | Motor detachment rate | $0.1 \text{ s}^{-1}$ | Estimated from ref.<sup>19</sup> |
| $k$ | Motor elasticity | $3 \text{ pN}/\mu\text{m}$ | Estimated from ref.<sup>55</sup> |
| $n_{\text{MT}}$ | Number of MTs in a nucleus | 10 | Measured here |
| $\langle R_{1,2} \rangle$ | Expected MT length | $0.8 \mu\text{m}$ | Ref.<sup>29,56</sup> |
| $D_{1,2}$ | MT diffusion constant | $0.003/R_{1,2}^3$ | Ref.<sup>29</sup> |
| $d_{\text{SPB}}$ | Distance between SPBs | $0.5 - 2.5 \mu\text{m}$ | Measured here |
| $R_C$ | Nuclear envelope radius | $1.5 \mu\text{m}$ | Ref.<sup>29,57</sup> |

### The model predicts that bundle formation is faster for small distances between the SPBs, large MT number and fast MT diffusion

To provide a quantitative measure that can be compared with experiments, we calculate the average bundling time, defined as the time required for MTs to form an antiparallel bundle (Methods). We found that the average bundling time is roughly 1-10 minutes for parameters in Table 1 and random initial conditions (Fig. 5a). The average bundling time increases as the SPB distance increases (Fig. 5a). To compare these results with experiments, we calculated the average spindle reassembly time for different distances between the SPBs by using the data from Fig. 1d, and found that our model reproduces the experimental measurements (Fig. 5a). There is a discrepancy at small SPB distances, possibly due to the smaller MT numbers at the onset of MT growth (Fig. 1e), which is the time window relevant for reassembly in this case. We conclude that the agreement between the model and experiments supports the hypotheses used to build our model.

We further explored the predictions of the model by varying the parameters of the model and calculating the resulting average bundling time (Fig. 5b). We found that an increase in MT number by a factor of 1.6 accelerates bundle formation by a factor of 3, whereas a decrease in MT number slows down this process. Similarly, MT diffusion affects bundle formation, though to a smaller extent (Fig. 5b). On the other hand, the number of motors and their stiffness has a minor contribution (Fig. 5b). Thus, bundling time depends mainly on MT number and diffusion.

**Figure 5.**
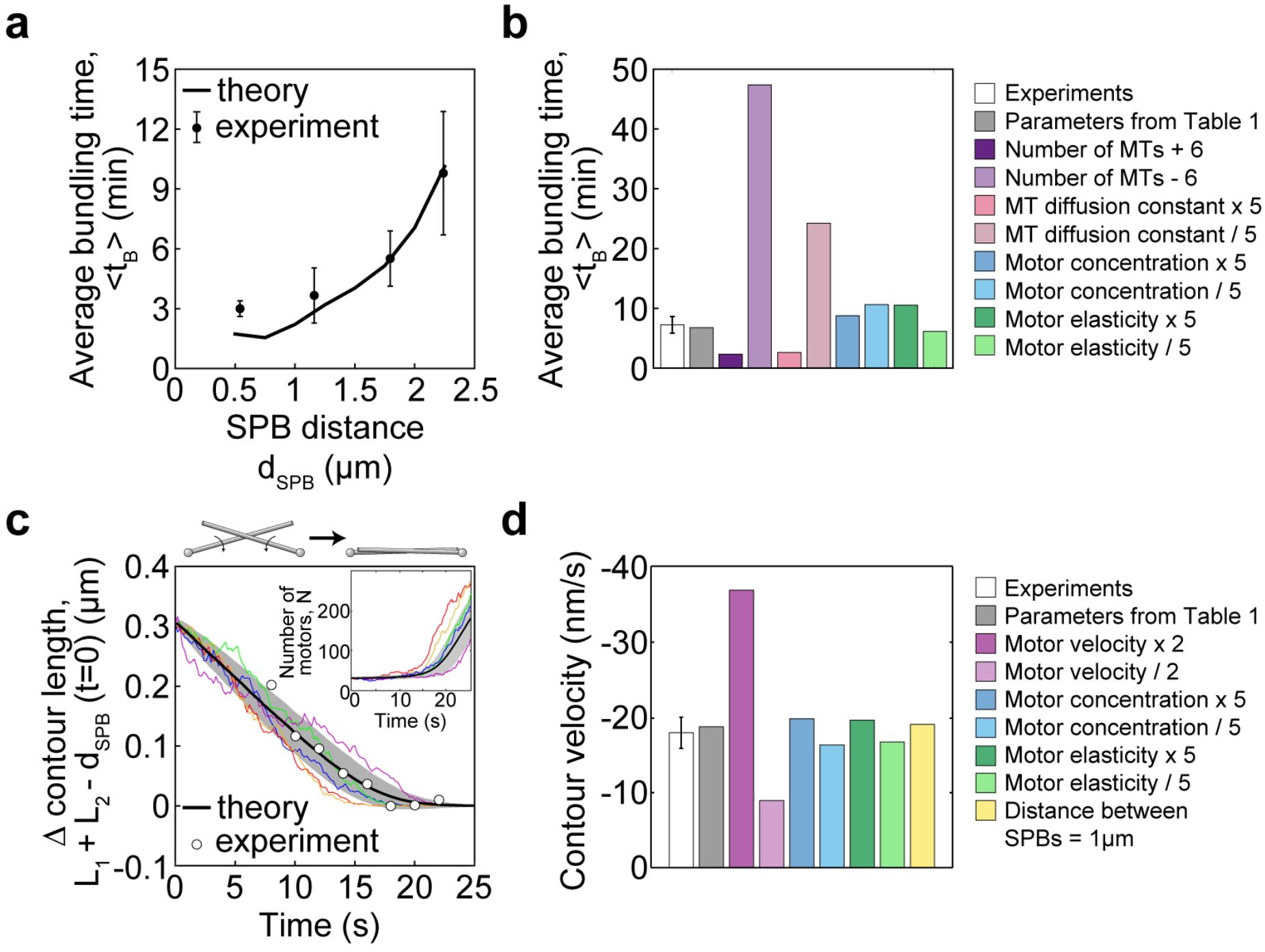
Bundling time and contour length. (**a**) Comparison of average bundling times in the theory and the experiment. The black line represents the average bundling time obtained from numerical simulations for different values of *d*_SPB_. The parameters used are shown in Table 1. The circles with error bars represent the average reassembly time measured in experiments for *d*_SPB_ < 2.5 *μm.* The average bundling time is defined 〈*t*_B_〉 = *t*_tot_/*n*_B_, where *t*_tot_ is the total time the MTs were observed or simulated and *n*_**B**_ is the number of bundling events in that time. The error bars are s.e.m. (**b**) Average bundling times in experiments (spindle reassembly time, white bar, error bar is s.e.m.) and in theory for different parameter values (colored bars). Deviations from parameter values used in Table 1 are denoted in the legend. The experimental value is for 1.5 *μm* < *d*_SPB_ < 2.5 *μm*, and the theoretical values are for *d*_SPB_ = 2 *μm.* The average bundling time is calculated as in (**a**). (**c**) Contour length difference (main graph) and the number of motors (inset) as a function of time. In the simulations, MTs start from a symmetric configuration in which the angle between them is 120°, the distance between the MTs is negligible, *y*_min_ ≈ 0 *μm.* We define the contour length as the sum of the lengths *L*_1_ and *L*_2_, which extend from the SPB to the point on the MT that is closest to the other MT, and the contour length difference is calculated as *L*_1_ + *L*_2_ − *d*_SPB_. The black line and the shaded area represent the average value and standard deviation, respectively, for all simulation results at a given time point. The colored lines are sample paths for simulations. The white dots represent the mean experimental values shown in Fig. 2d, with their origin on the time axis shifted by the average bundling time of the simulation runs. Simulations are performed with the parameters form Table 1, *d*_SPB_ = 2 *μm*, *R*_1,2_ = 2 and *n*_MT_ = 2. (**d**) Contour velocity in experiments (white bar; error bar is s.e.m.) and in simulations for different parameter values (colored bars). Deviations from parameter values in Table 1 and *d*_SPB_ = 2 *μm* are denoted in the legend. MT lengths are *R*_1,2_ = 2 *μm*, except for the last bar, where *R*_1,2_ = 1 *μm.* Initial configurations are the same as in (**c**).

### Minus end directed motors align the MTs into an antiparallel bundle at a constant velocity

In our experiments, we found that the total contour length of MTs decreased during their rotation into antiparallel alignment. To help us interpret these results, we used the model to explore the change in MT contour length. Here we start our calculations from the configuration in which the MTs are in contact (Fig. 5c is for a symmetric and Supplementary Fig. S3a for an asymmetric initial configuration). We find that the contour length difference decreases towards zero and remains constant afterwards (Fig. 5c). This process is driven by motors, which initially accumulate slowly, whereas the accumulation accelerates as the MTs approach the aligned configuration (Fig. 5c, inset). The contour length decreases linearly at a velocity close to 2*v*_0_, twofold the velocity of motors along each MT. The value of the parameter *v*_0_ was chosen to reproduce the experimentally measured contour velocity (Table 1 and Methods). To explore the predictions of the model, we varied the parameters and found that the contour velocity is proportional to the motor velocity, whereas the number of motors, their stiffness, and the SPB distance have a minor contribution (Fig. 5d and Supplementary Fig. S3b; other parameters are investigated in Supplementary Fig. S3c). Interestingly, the prediction that the contour velocity does not depend on the SPB distance implies that this velocity is robust to changes in the geometry of the system. To test this prediction, we divided the cells into two groups, those with the SPB distance smaller or larger than 1.85 μm. Indeed, we found that the contour velocity was not different between these groups (−18±5 nm/s and −24±6 nm/s for the cells with SPB distance of 1.69±0.03 μm and 2.3±0.2 μm, respectively; n=14; p=0.5 from a t-test for velocities). Taken together, our results suggest that the minus end directed motors align the MTs into an antiparallel bundle and allow us to identify their velocity.

Our model together with experiments shows that the search times are severalfold longer than the duration of MT aligning. Thus, the average bundling time, which includes both search and aligning, describes predominantly the time scale of the search process, which is on the order of minutes. The change of contour length captures only the time scale of the aligning step, which is on the order of seconds. Our model, which is based on the known properties of MTs and motors, explains the entire process of antiparallel bundle formation covering different time scales.

## DISCUSSION

### Microtubule pivoting around the spindle pole facilitates their encounter

By using a spindle reassembly assay, we found that the formation of an antiparallel bundle occurs in two steps, search and aligning (Fig. 1c). During the search step, MTs extending from the opposite SPBs rotate around the SPB, which helps the MTs to find each other. MT rotation during the search step is passive angular diffusion, which is thermally driven and does not require ATP (ref.^29^). Previous computer simulations of spindle assembly in fission yeast indicate that a decreased MT rotation results in fewer MTs in the bundle connecting the two SPBs and shorter spindles^32^. Thus, previous work and our model together with experiments show that rotation of MTs is required for the process in which they search for each other to form an antiparallel bundle.

MT rotation has been observed before in fission yeast during mitosis and meiosis^29,56^, in budding yeast^58^, and in *Drosophila* S2 cells^59^. In general, pivoting helps the MTs as they search for targets such as kinetochores^29,56,60^, cortical anchors *in vivo*^58^ and *in vitro*^61^, or other MTs (ref.^30,59^). This motion allows MTs to swipe through space, which increases the explored volume and makes the search process more efficient^2,62^.

### Dynamics of the microtubule contour length reveals that minus end directed motors align the microtubules into antiparallel bundles

During the second step of bundle formation, MTs extending from the opposite SPBs rotate towards the pole-to-pole axis to form an antiparallel configuration. This rotation occurs after the MTs have established contact at an arbitrary angle. Whereas MT rotation is random before the contact, it becomes directed as they pivot towards the antiparallel alignment after the contact. By using a spindle reassembly assay to increase the distance between the spindle poles, we were able to observe and quantify this directed rotation. MT contact at oblique angles has been observed in electron micrographs of cells in early mitosis^28^, thus we propose that these MTs rotate to become aligned and form an interpolar bundle in unperturbed cells. Moreover, our results may be relevant for higher eukaryotic cells, where the majority of antiparallel bundles form when the centrosomes are apart^63^.

During MT alignment, MTs slide sideways along each other like a skater on a handrail. We introduce the contour length of MTs as a measure of activity of motors that drive MT sliding. This measure provides information about motor directionality and velocity. Our finding that the contour length of MTs decreases during their rotation, together with our theory, implies that the motors accumulated at the contact site walk towards the minus ends. Our measurement that the contour length decreases at a velocity of roughly 20 nm/s indicates that the motors walk at this velocity. This reasoning holds for fission yeast spindles where poleward flux is absent^40^. In the case of a tetrameric motor, the motor walks along each MT at a half of that velocity.

The velocity measured here is similar to the velocity of the minus end directed motility of kinesin-5 and kinesin-14 motors, but smaller than dynein velocity, in yeasts. In gliding assays *in vitro*, the kinesin-5 Cut7 from fission yeast moves at a velocity of 30 nm/s (ref.^23^), and Cin8 from budding yeast at 30-50 nm/s (ref.^19^). The kinesin-14 Pkl1 from fission yeast moves at a velocity of 33 nm/s in a gliding assay^64^, while the truncated Kar3 from budding yeast moves at 20 nm/s in a gliding assay^65^ and the full-length Kar3-Cik1 at 45 nm/s in a stepping assay^31^. Finally, cortically anchored dynein in fission yeast moves the nucleus at a velocity of 100 nm/s (ref.^66^), and single dyneins move at 140 nm/s along the MT (ref.^67^), whereas full-length purified budding yeast dynein glides MTs in vitro at 90 nm/s (ref.^68^).

### Molecular players involved in resolving oblique microtubule contacts

We propose that Cut7, a kinesin-5 family member, plays a role in MT rotation into antiparallel alignment, based on the literature and our experiments. Cut7 is essential for spindle formation^13^, unlike the other 8 kinesins of *S. pombe*^42–47^, dynein^48^ and the non-motor crosslinker Ase1/PRC1 (ref.^49,50^). The key role of Cut7 in spindle formation was revealed by inactivation of Cut7 in temperature-sensitive mutants, which resulted in cells with monopolar spindles^13^. Similarly, Cut7 inactivation during metaphase leads to spindle collapse into an aster pattern^47^. Electron micrographs of monopolar spindles produced by Cut7 inactivation showed that MTs extending from the two SPBs are roughly parallel, suggesting that Cut7 is required for MT interdigitation^69^. Our experiments showing accumulation of Cut7 at the site of initial MT contact, together with low efficiency of spindle reassembly in a temperature-sensitive *cut7.24*^ts^ mutant, are consistent with a role of Cut7 in the formation of antiparallel MT bundles starting from oblique MT interactions.

Other motors, such as the minus-end directed kinesin-14 motors Pkl1 and Klp2 (ref.^42,43^) and dynein^48^ might contribute to MT rotation into antiparallel alignment. Yet, we found that spindles were able to reassemble in cells lacking Pkl1 or Klp2, which is consistent with previous observations that spindles can reassemble in cells lacking Pkl1, Klp2 or dynein after MT depolymerization in kinetochore capture assays^38,70^. Additionally, cells lacking any of these three motors or even all three of them are able to form spindles under normal conditions^43,48^. Thus, Pkl1, Klp2 and dynein are not crucial for the resolution of oblique MT contacts during spindle assembly.

Finally, it is possible that MT rotation into antiparallel alignment occurs without motor activity. However, based on our measurements of contour length together with the theoretical results, we favor the interpretation that the action of minus end directed motors drives MT alignment. Our experiments on Cut7 together with the previously observed minus end directed motility of Cut7 (ref.^23^) suggest a role of Cut7 in this process.

### Change in the direction of forces after microtubule alignment

While our model together with experiments indicates that motors walk towards the MT minus end to align MTs, this minus end directed motility is expected to shorten the bundle after it is formed. Yet, we observed that the spindle remained at a constant length or elongated slowly upon reassembly, which is indicative of forces acting in the opposite direction. Thus, our work suggests that there is a switch in the direction of forces during spindle reassembly, and the underlying mechanisms are currently unknown. It may be that forces generated by other molecular players, which push the spindle poles apart, start to dominate over the minus end directed motors. These forces may be generated by plus end directed motors such as kinesin-6/Klp9 (ref.^47^), or by pushing forces generated by the interaction of growing MT plus ends with the opposite SPB (ref.^33,34^). Alternatively, the motors that align the MTs into the antiparallel bundle may change their direction of motion after the bundle is formed. Crowding on the MT has been suggested to convert the Cut7 motor from minus end directed to plus end directed stepping^20^. Similarly, single Cin8 motors from budding yeast were shown to move towards the minus end on individual MTs, but they switch to plus end directed motility when working in a group of motors on antiparallel MTs^19,21^. Exciting new experimental and theoretical investigations await in this field to reveal how the regulation of motor protein activity and MT dynamics govern spindle formation and function.

## ACKNOWLEDGMENTS

We thank Phong Tran, Jonathan Millar, and Yeast Genetic Resource Center for strains and plasmids; Falk Elsner from the Electronics Service of MPI-CBG for building the thermoelectric device; Britta Schroth-Diez and Jan Peychl from the Light Microscopy Facility of MPI-CBG for help with microscopy; Ivana Šarić for help with image processing and editing the figures; Stefan Diez, Jan Brugues, and members of Tolić and Pavin groups for discussions and advice. This work was funded by the Max Planck Society and Unity through Knowledge Fund (UKF, project 18/15 granted to N.P. and I.M.T.). We also acknowledge support from the German Research Foundation (DFG, project TO 564/7-1 granted to I.M.T. and N.P.), QuantiXLie Centre of Excellence, a project cofinanced by the Croatian Government and European Union through the European Regional Development Fund - the Competitiveness and Cohesion Operational Programme (Grant KK.01.1.1.01.0004, element leader N.P.), and the European Research Council (ERC Consolidator Grant, GA Number 647077, granted to I.M.T.).

## AUTHOR CONTRIBUTIONS

L.W. carried out experiments; L.W., I.B., M.P., and I.M.T. analyzed data; I.B., M.P. and N.P. developed the theoretical model; I.B. and M.P. solved the model; I.M.T. and N.P. conceived the project and supervised experiments and theory, respectively. I.M.T. and N.P. wrote the paper with input from all authors.

## METHODS

### Strains and sample preparation

The strains (Table S1) were obtained by crossing, followed by random spore analysis ^71^. The cells were grown on Yeast Extract with supplements (YES) medium agar plates at 25°C ^71^. A loopfull of cells was further cultured in liquid YES medium in a shaking incubator (ISF-1-W, Kuehner Shaker, Birsfelden, Switzerland) at 25°C for 2-3 hours. For the strains CF.391, I1_2_10, KI013, and LW042, 3 mM hydroxyurea (Sigma-Aldrich) was added to liquid YES medium in order to obtain lengthy cells with normal MT dynamics ^72,73^ and with lengthy spindles, the cells were kept for 11-14 hours in the shaking incubator at 25°C, and subsequently the liquid culture was diluted with liquid YES medium at a ratio 1:3. The wall of a 35 mm (No1.5) culture dish (MatTek Corporation, Ashland, MA, USA) was cut to 2 mm height. The original coverslip was removed from the dish bottom and the remaining dish was soaked in 70% ethanol overnight. Cover slips (Corning 22 mm x 22 mm, Sigma-Aldrich) were washed in 2-propanol and attached to the pre-washed culture dish with transparent nail polish. The dish was coated with lectin (L2380, Sigma-Aldrich, St Louis, MO, USA) 30 minutes prior to usage. 200 μl of liquid culture was placed on the pretreated culture dish for 25 minutes for sedimentation. The cells were washed 3 times with 200 μl of YES medium. The dish was closed with a cover slip (Corning, Inc.) to prevent the sample from drying out.

### Microtubule depolymerization by a custom designed thermoelectric device

To quickly depolymerize metaphase spindles, a thermoelectric device based on a Peltier element was designed and tested with an independent type K thermocouple (Omega Engineering, Deckenpfronn, Germany) and a Fluke 50 Serie II Thermometer (Fluke Corporation, Everett, WA, USA). Prepared samples were loaded onto the microscopy stand and a pre-cooled thermoregulation at 15°C, to slow down mitosis and thus facilitate our search for a field of view with a high number of cells in metaphase (Supplementary Fig. S1a). Following image acquisition, the temperature was set to 0°C. Once set to 0°C, the temperature dropped to 1°C within 60 seconds inside the sample and was maintained typically for 15 minutes. To ensure a constant temperature, the objective was lowered to at least 2 cm away from the sample dish. Subsequently, the objective was returned to the initial position, the same field of view was placed into focus, and image acquisition was initiated. Within 20 seconds of acquisition, the temperature was set to 24°C. The temperature inside the sample reached this value within 30 seconds. Because of the change in temperature, the sample was manually refocused during the acquisition. Once the temperature in the sample was stabilized at 24°C, the focus remained constant. In experiments with *cut7.24*^ts^ cells, the final temperature was set to 37°C instead of 24°C to inactivate Cut7. Live-cell imaging was performed for 10 minutes.

### Time-lapse live cell imaging

Live images were taken using an Andor Revolution Spinning Disk System (Andor Technology plc., Belfast, United Kingdom), consisting of a Yokogawa CSU10 spinning disk scan head (Yokogawa Electric Corporation, Tokyo, Japan) with a 405/488/568/647 Yokogawa dichroic beamsplitter (Semrock, Inc., Rochester, NY, USA). The scan head was connected to an Olympus IX71 inverted microscope (Olympus, Tokyo, Japan) equipped with a fast piezo objective z-positioner (PIFOC, Physik Instrumente GmbH & Co. K.G., Karlsruhe, Germany) and an Olympus UPlanSApo 100x/1.4 NA oil objective (Olympus, Tokyo, Japan). For cells expressing GFP and tdTomato, we performed sequential imaging (2 second time interval between each image pair) or simultaneous acquisition (1 second time interval between images) using a DualView image-splitter (Optical Insights, Photometrics, Tucson, AZ, USA). Cells expressing only GFP (Mal3-GFP) were imaged with 250 ms time interval. Exposure time was 20 ms. For excitation, a Sapphire 488 nm solid-state laser (75 mW; Coherent, Inc., Santa Clara, CA, USA) and a Jive 561 nm solid-state laser (75 mW; Cobolt, Stockholm, Sweden) were used for GFP and tdTomato, respectively. Laser intensity was controlled using the acousto-optic tunable filter inside the Andor Revolution Laser Combiner (ALC, Andor Technology plc., Belfast, UK). For sequential imaging, emission wavelength was selected using respective emission filters BL 525/30 (Semrock, Inc., Rochester, NY, USA) and ET 605/70 (Chroma, Bellows Falls, VT, USA) mounted in a fast, motorized filter wheel (Lambda-10B, Sutter Instrument Company, Novato, CA, USA). The microscope was equipped with an iXon EM+ DU-897 BV back-illuminated Electron Multiplying Charge Coupled Device (EMCCD, Andor Technology plc., Belfast, UK), cooled to - 80°C, electron multiplication gain 300. The resulting xy-pixel size in the images was 168 nm. The system was controlled by Andor iQ software version 2.9 (Andor Technology plc., Belfast, UK). For short-term acquisitions (10-20 seconds), sequential time-lapse z-stacks (2-second time interval between each image pair) of 13 optical sections with 0.5-μm z-spacing was performed using a DualView image-splitter (Optical Insights, Photometrics). For main acquisitions (10-minutes), time-lapse z-stacks of 13 optical sections with 0.5-μm z-spacing were taken every 2 seconds with exposure times of 0.06 and 0.08 seconds. In the case of main acquisitions (10-minutes) of strain LW042, sequential time-lapse z-stacks (5-second time interval between each image pair) of 13 optical sections with 0.5-μm z-spacing was performed, with exposure times of 0.08 and 0.1 seconds for GFP and mCherry, respectively.

### Theoretical model

#### Orientations of the MTs

We model the MTs as two thin, rigid rods of fixed length *R*_1_ and *R*_2_ (here and in the rest of this text, indices 1 and 2 represent the first and the second MT, respectively), each with one end freely joint (pinned, but not clamped) at the respective SPB. Their orientations are represented by unit vectors **r̂**_1,2_ (Fig. 4a in the main text), and the SPBs are positioned at the origin and at **d**_SPB_ = *d*_SPB_**ẑ**, with **ẑ** being the unit vector in the direction of the Cartesian z-axis. The MTs pivot around their respective SPB with the angular velocities *ω*_12_. The orientations of MTs change in time, *t*, as

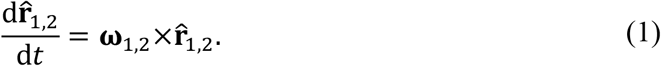

In the overdamped limit, the angular friction experienced by the MTs is balanced by the total torque,

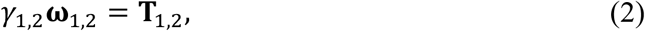

where *γ*_1,2_ is the angular friction coefficient of the MTs. The total torque consists of two contributions, **T**_1,2_ = τ_l,2_ + *σ*_1,2_[**r̂**_1,2_×**η**_1,2_(*t*)], where the first term is the deterministic torque, τ_1,2_, caused by the attractive forces exerted by the motors attached to the MTs and the second term is the stochastic term describing the noise. In our model, the noise is thermal, so its intensity is calculated from the equipartition theorem as 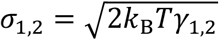 with *k*_B_*T* being the Boltzmann constant multiplied by the temperature. The 3-dimensional random vector **η**_1,2_ has components that are normally distributed with zero mean and unitary variance. The noise is uncorrelated in time and its components are independent, 〈η_*i*_(*t*),η_*j*_(*t*′)〉 = *δ*(*t* − *t*′)*δ_ij_*, with δ(*t* − *t*′) being the Dirac delta function and *δ*_*ij*_ is the Kronecker delta function. Using these definitions and equation (2), we obtain the equations for the angular velocities of the MTs,

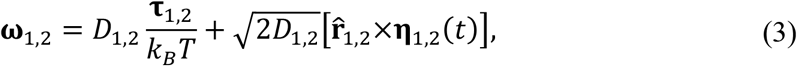

where *D*_1,2_ = *k*_B_*T*/γ_l,2_ denotes the angular diffusion coefficient of the MTs.

#### Forces, torques and motor movement

The torques in equation (3) depend on the distributions of the motors which are attached at a given time, 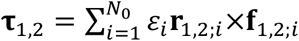. Here, the motor state has a value *ε*_*i*_ = 1 for a motor attached to both MTs and *ε*_*i*_ = 0 otherwise. The motors are labeled with indices *i* = {1,…, *N*_0_}, where *N*_0_ is the total number of motors.

A motor is modeled as a spring with ends that can move when attached to two different MTs. For a motor with its ends attached at the distances *r*_1,2;*i*_ from the respective SPBs along the MTs, the vector describing the elongation of its spring is **y**_*i*_ = **r**_2;*i*_ + **d**_spB_ − *r*_1;*i*_, where the positions of the motor ends with relative to the respective SPBs are **r**_1,2;*i*_ = *r*_1,2;*i*_**r̂**_1,2_ (Fig. 4a). The springs are Hookean with zero rest length, so the force they exert on the MTs is

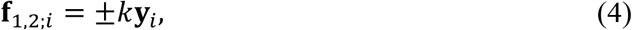

where *k* is the spring stiffness. The motor moves along both MTs and the velocities of the motor ends depend on the load experienced by them. We use a linear relationship between force and velocity, thus the positions of the motor ends change as

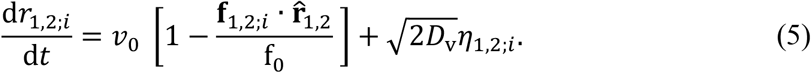

Here, the motor velocity, *ν*_0_, is the velocity of a motor head at zero load, the stall force is *f*_0_, and the scalar product is the component of the force parallel with the MT. The motor velocity fluctuation, *D*_v_, describes the natural stochasticity in the movement of the motor head.

#### Motor attachment and detachment

In order to determine the distribution of the motors, we start by modeling a single motor, dropping the indices *i* to simplify the notation, and then generalizing to the total number of motors, *N*_0_. We introduce the motor state process *ε* ≡ *ε*(*t*). Initially, at *t* = 0, the motor can be either in the state *ε* = 0 or *ε* = 1. Here, we construct the motor state process for the case *ε*(0) = 0. Motor attachment is a jump process with the rate *k*_on_(**y**), which depends on the motor elongation and the MT orientations. Both the motor elongation and the MT orientations change stochastically in time, so the attachment process is defined by assigning each small time interval [*t*, *t* + Δ] an attachment probability *p*_a_(Δ*t*) = *k*_on_(**y**)Δ*t* and taking the limit Δt→0. Because the free motor experiences thermal fluctuations, the elongation at each time interval is a random variable that has a Boltzmann distribution. The time point at which the motor state changes to attached, *ε* = 1, is denoted *t_1_*.

Once the motor is attached, its detachment is a Poisson process with a constant rate *k*_off_, thus the time interval in which the motor stays attached is an exponentially distributed random variable, (*t*_2_ − *t*_1_)~ Exp(*k*_off_), where *t*_2_ denotes the time point at which the motor state changes back to *ε* = 0. The next attachment times *t*_3_, *t*_5_,… are defined analogously to *t*_1_ and the detachment times *t*_4_, *t*_6_,… analogously to *t*_2_. Therefore, a sequence of random time points representing attachment and detachment events, 0 < *t*_1_ < *t*_2_ … < *t*_2*j*−1_ < *t*_2*j*_ …, defines the motor state process, *ε*(*t*). Here, the *j*-th attachment event occurs at odd-indexed time points *t*_2*j*−1_, while the *j-th* detachment event occurs at even-indexed time points *t*_2*j*_. In order to calculate the motor’s contribution to the total torque in equation (3), we calculate the position of each motor head by solving equation (5) in the time intervals *t*_2j−1_ < *t* < *t*_2*j*_, with the initial conditions for each interval *r*_1,2_(*t*_2*j*−1_) calculated from **y**(*t*_2*j*−1_). For a motor that starts attached to the MTs at initial distances *r*_1,2_(0), we keep the index parity convention: the first event that occurs is a detachment event at a time point we denote *t*_2_, then an attachment event at *t*_3_, and so on as described above. In this case, the time sequence of attachment and detachment events is 0 < *t*_2_ < *t*_3_ …

The generalization to *N*_0_ motors is straightforward. The sequence of attachment times, {*t*_2*j*−1;*i*_}, and detachment times, {*t*_2*j;i*_}, is calculated for each motor separately. At a given time, there are 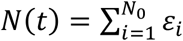 motors that are attached and contribute to the torque with coordinates, *r*_12;!_, given by equation (5). One has to keep in mind that, while the motors attach, detach and move independent of each other, the dependence of the attachment rate, *k*_on_(**y**_i_), and the projection of the force, **f**_1,2;*i*_**·r̂**_1,2_, in equation (5) on MT geometry, which in turn depends on the positions of attached motors (equations (1-3)) introduces coupling between them.

### Solutions of the model

To obtain the time course of the MT orientations, we parameterize the orientation of the MT given by the unit vector by **r̂**_1,2_(*θ*_1,2_, **ø**_1,2_) = (sin *θ*_1,2_ cos *ø*_1,2_, sin *θ*_1,2_ sin *ø*_1,2_, cos θ_1,2_), where *θ*_1,2_ and *φ*_1,2_ denote the polar and azimuthal angle, respectively. The equations of motion for the angles are obtained by expressing equation (3) in spherical coordinates,

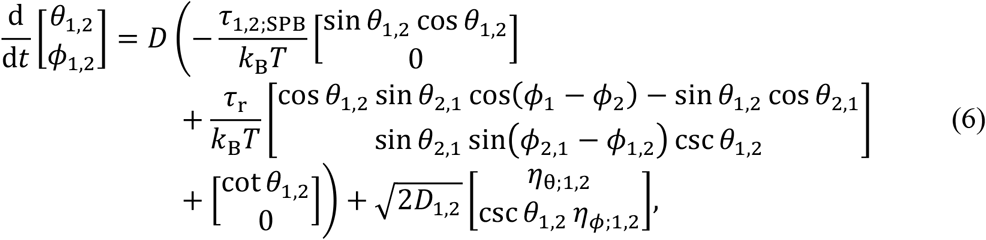

The additional deterministic term (usually called spurious drift), *D* cot*θ*_1,2_, appearing in the equation for the polar angle and the factor csc *θ*_1,2_ in the noise for the azimuthal angle are the results of coordinate transformations. For a derivation, see ref.^30^. The torque magnitudes are 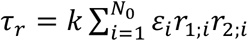 and 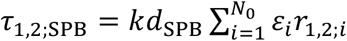. The second sum has terms linear in *r*_1,2;*i*_, meaning that it can be evaluated without knowing the exact coordinate of each motor, only the average position r̄_1,2_ and their number, *N*, i.e. 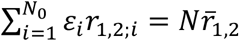. Because the motors are much smaller than the MT, they only accumulate where the MTs are close to each other, so we can approximate the first sum as τ_l,2;*r*_ ≈ *kL*_1,2_*Nr̄*_1,2_, where *L*_1,2_ are the points on the respective MTs that are closest to each other. Because of this approximation, the torque in equation (6) depends only on the number of attached motors and their average positions.

In the limit in which the number of motors in the nucleoplasm, *N*_0_, is large, the probability of finding *N* motors attached to both MTs, *p*_*N*_, is calculated using the master equation

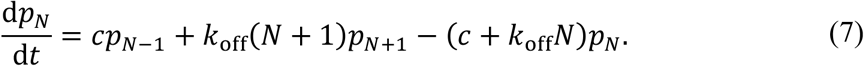

The effective number of motors in the nucleoplasm is given by

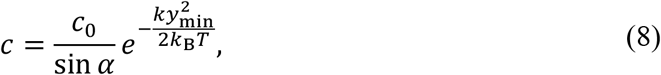

where the constant *c*_0_ ∝ *N*_0_ is termed the motor concentration parameter, *α* is the angle between the MTs and *y*_min_ = *y*(*L*_1_, *L*_2_) is the minimal distance between the MTs. Finally, for *N* ≫ 1, *N(t)* can be considered a continuous variable and equation (7) can be approximated with a Langevin equation for the number of motors,

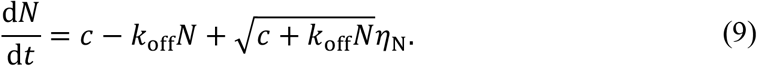

This equation is derived by calculating the expected value and variance of the number of motors, 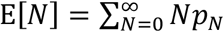 and the variance, 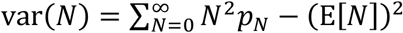 from equation (7).

In order to calculate the average positions of the motors, it is convenient to introduce the auxiliary coordinates *u* = (*r*_1_ + *r*_2_)/2 and *w* = (*r*_1_ − *r*_2_)/2, and using them in equations (5), yielding two independent equations. The torques can be calculated by transforming back using the expressions *r̄*_1,2_ = *ū* ± *w̅*, where *ū* and *w̅* are the average values. Aside from motor movement along MTs, motor attachment and detachment affects the average coordinate of the motors. By considering the jump processes in the continuous limit and combining with the continuous contributions expressed in equation (5), we obtain two independent Langevin equations for the average coordinates of the motors,

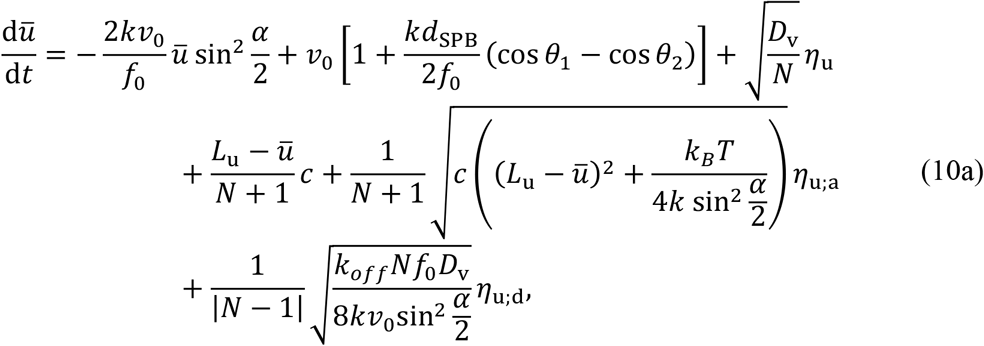

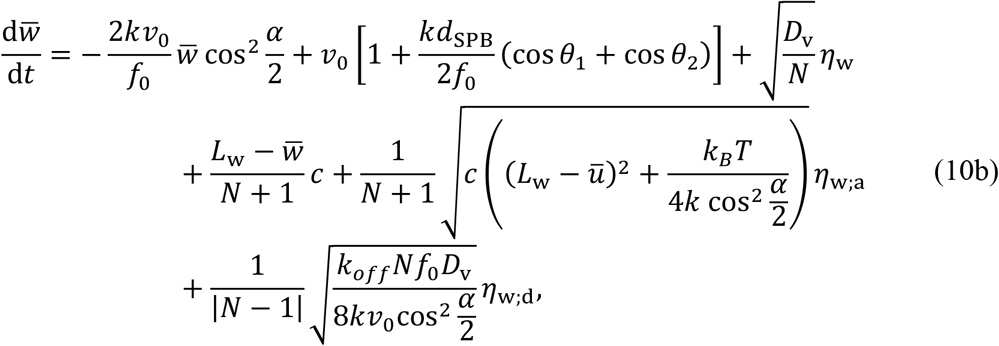

where *L*_u,w_= (*L*_1_ ± *L*_2_)/2. The first three terms on the left in either equation are due to motor movement, the next two terms are due to motor attachment and the last term is due to motor detachment. The detailed derivations of equations (9) and (10) are given in [Prelogović et al., unpublished, available on request].

Using the average coordinates given in equations (10a,b) and the number of attached motors given by equation (9), we calculate the torque components in equation (6), which then yield the time course of the MT orientations, thus solving the model.

### Choice of parameter values

Our model has 11 parameters, 1 of which is free (motor concentration parameter *k*_on_*c*_nuc_). We have 5 parameters related to motors, which we estimated based on previous *in vitro* measurements for kinesin-5. The movement of motors is described by their velocity at zero load, *v*_0_ = − 0.01 μm/s, which we estimated as half the measured contour velocity (roughly −0.02 *μm/s*), given that the tetrameric motors walk along each MT with half of this velocity. This velocity is 3 times smaller than velocity measured from *in vitro* motility assays for Cut7 (ref.^23^), 2 times smaller than velocity for Kip1 purified from *S. cerevisae* from *in vitro* experiments, at high ionic strength conditions in ref.^22^, and 6 time smaller than velocity for purified Cin8 in budding yeast measured in ref.^19,21^. The motor velocity dispersion, *D*_v_ = 4×10^−4^μm^2^/s, is estimated based on theory (*D*_v_ = *rv*_0_*d*/2, where *d* = 36 *nm* is the kinesin step size and *r* ≈ 0.39 is the randomness observed in optical tweezer experiments^53^). For stall force, we used *f*_0_ = −1.5 pN measured for Cin8 from budding yeast^54^, which is consistent with the stall force estimated for Xenopus kinesin-5 (ref.^74^), but five times smaller than reported in human kinesin-5 dimeric construct^75^. The off rate 0.1 *s*^−1^ of motor detachment is estimated based on/from the dwell time for Cin8 (ref.^19^), which is similar to the value used in ref.^32^, and two times smaller than off rate for Kip1 in (ref.^22^, estimated as the motor velocity divided by its run length). Value for the motor concentration parameter is roughly estimated so that there are 30 motors attached when the MTs are in contact and the angle between them is 120°, *k*_on_*c*_nuc_ = 1.3 *s*^−1^μm^−1^. The motor is described as a Hookean spring with zero rest length whose elasticity is calculated so that the expected motor length in thermal equilibrium matches the length of the crossbridge, 53 *nm*, Fig. 3c in ref.^55^, which is similar to the motor rod length measured in ref.^76,77^ and equal to the one used in ref.^32^.

For the MTs, we have 2 parameters: diffusion constant *D*_1,2_ and MT length *R*_1,2_. We calculate the diffusion constant as *D ∝ R^−3^* (ref.^78^) using fitting results from ref.^29^, which is consistent with ref.^30^. For MT lengths, we assume they follow an exponential distribution. The expected value of measured MT lengths is 1.5 μm (ref.^29^). We assume that the distribution of MT lengths is exponential^56^, but measurements only take into account MTs longer than 0.7 μm, so the real distribution consistent with the measurements is *R~* Exp(1/0.8 μm^−1^), yielding the expected value of *R*_1,2_ = 0.8 μm. In our experiments, the average number of visible MTs on each SPB is 4, which means there are on average 10 MTs in a real cell, using the above exponential distribution. Assuming there is a roughly equal number of MTs at each SPB, this implies that there are around 25 possible combinations of MTs that can form a bundle, and the real bundling time is the fastest bundling time out of these combinations. We varied the distance between the SPB in the range to match the variability among the cells in our experiments. The nucleus is approximated as a sphere with radius *R*_*c*_ = 1.5 μm, a value that is estimated from the nuclear volume^57^.

### Numerical simulations

In order to obtain the average bundling times and contour velocities, we solved the system of equations (6), (11) and (20a,b) numerically by simulating the sample paths. The simulations were performed using an Euler-Maruyama scheme for solving stochastic differential equations, with a reflective boundary condition representing the nuclear envelope (the envelope was assumed to be a hard spherical shell with both SPBs embedded in it). The simulations we performed slightly differently for obtaining bundling times and contour velocities.

For calculating the average bundling time, we simulated 3000 sample paths per value of *d*_SPB_ for MT angles, which lasted for *t*_max_ = 10 *min* or until the bundling angle between MTs of *α*_b_ = 3.05 was reached. The MT lengths were generated randomly from an exponential distribution discussed under Table 1. The initial condition for the angles was randomly generated so that the initial polar angles have a sinusoidal distribution and the azimuthal angles have a uniform distribution. If the randomly generated initial orientation would place the MT outside of the nucleus, it would be rejected and generated again. The last time of each run was recorded. In order to represent the fact that there are many MTs on each SPB, the run times were randomly organized into sets of (*n*_MT_/2)^2^, and only the fastest time would represent a single data point. This was done 10000 times.

For simulating the contour length in time, we set the initial condition for the angles in radians (*θ*_1_, *θ*_2_, *ø*_1_, *ø*_2_)|_*t*=o_ = (0.5, 2.6, 0.001,0.0001), and fixed the MT length to be the same as the SPB distance, *R*_1,2_ = *d*_SPB_. We then performed 1000 runs for parameter values shown in Table 1 to obtain the sample paths. At each time point, the mean and standard deviation were calculated for all simulation runs (see Fig. 5c). The contour velocities shown in Fig. 5d were calculated by repeating the simulations for each value of the parameters of interest and performing a linear fit on the simulated data in the time interval 1 *s* < *t* < 5 *s*. The experimental velocity was obtained by performing the linear fit on all of the experimental data before the bundling time.

### Quantification and statistical analysis

*Maximum-intensity projections* were calculated with ImageJ (National Institutes of Health, Bethesda, MD, USA) using the plug-in Grouped Z Projector under Stacks selection. The color-merge images were obtained by overlay of projections in green and red channels using Merge Channels under Color selection.

#### Tracking of MTs and SPBs

The position of MT tips was manually tracked by using Manual Tracking plug-in under Tracking selection. MT tracking was performed on MTs longer than 0.5 μm that appeared before spindle reassembly with traces longer than 1 minute. In the strains in which SPBs were fluorescently labeled, SPBs were tracked by specialized tracking software ^79^, whereas in other strains the position of the SPBs was estimated based on the end of MT signal.

*The average spindle reassembly time* was calculated as the total time of spindle reassembly over all cells (for non-assembled spindles this time is equal to the duration of imaging, i.e., 10 minutes) divided by the number of reassembly events. The error (s.e.m.) was calculated as the average reassembly time divided by the square root of the number of reassembly events.

*The total contour length of MTs* was measured in 14 out of 21 cells in which MTs interacted at an oblique angle. In the remaining 7 cells, the contour length was not measured because the point of MT contact was not clearly visible in all time frames. We measured the contour length up to 10 seconds before and 4 seconds after the first antiparallel bundle was formed (time *t*=0). We measured the contour length using Multi-point tool in Fiji. In each time frame we measured the coordinates of three points: at one SPB, at the point of MT interaction and at the other SPB (Supplementary Fig. S1e). From the coordinates of these points, we calculated the angle between MTs, the contour length, and the distance between SPBs.

*Signal intensity profiles* of mCherry-tubulin and Cut7-3GFP were measured by using a Freehand Line tool in Fiji. The line was drawn starting from in the cytoplasm, passing though one SPB, following the MT contour via the contact point, passing through the other SPB and ending in the cytoplasm. A new line was drawn on each time frame in the channel for mCherry-tubulin.

Data are presented as mean±s.e.m. The error (s.e.m.) on *proportion data* was calculated as 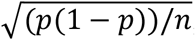 where *n* is the sample size, and *p* is the number of events divided by *n*. Data analysis was performed using Excel (Microsoft Corporation) and custom scripts written in MATLAB (Mathworks). Figures were assembled in Adobe Illustrator (Adobe Systems).

### Data availability

The authors declare that all data supporting the findings of this study are available within the article and its supplementary information files or from the corresponding authors upon reasonable request.

### Code availability

The code used in this study is available from the corresponding authors upon reasonable request.

